# Ultrasensitive molecular controllers for quasi-integral feedback

**DOI:** 10.1101/413914

**Authors:** Christian Cuba Samaniego, Elisa Franco

## Abstract

Feedback control has enabled the success of automated technologies by mitigating the effects of variability, unknown disturbances, and noise. Similarly, feedback loops in biology reduce the impact of noise and help shape kinetic responses, but it is still unclear how to rationally design molecular controllers that approach the performance of controllers in traditional engineering applications, in particular the performance of integral controllers. Here, we describe a strategy to build molecular quasi-integral controllers by following two design principles: (1) a highly ultrasensitive response, which guarantees a small steady-state error, and (2) a tunable ultrasensitivity threshold, which determines the system equilibrium point (reference). We describe a molecular reaction network, which we name Brink motif, that satisfies these requirements by combining sequestration and an activation/deactivation cycle. We show that if ultrasensitivity conditions are satisfied, this motif operates as a quasi-integral controller and promotes homeostatic behavior of the closed-loop system (robust tracking of the input reference while rejecting disturbances). We propose potential biological implementations of Brink controllers and we illustrate different example applications with computational models.

## 1 Introduction

Feedback control enables the operation of most automated systems, from laptops to self-driving cars. Feedback works by reducing the discrepancy between the actual and desired behavior of the process to be automated, via a rationally designed controller (Fig. 1A). For example, a car cruise control system measures the speed of the vehicle, compares it to the set-point speed, and programs the fuel injection to reduce the error (fuel injection is increased if the speed is lower than the set-point, or decreased if the speed is higher). This architecture provides two key advantages: 1) Robustness: the process can maintain its set-point in the presence of disturbances (for example, changes in the slope of the road) or uncertainty in the process parameters (for example, the weight of the vehicle and passengers may not be known); 2) Response design: the speed and steady-state of the process response can be tuned easily (the time it takes to reach the desired speed can be reduced with a more aggressive controller). Robustness and tuning of the response are obtained exclusively by designing the appropriate controller, without the need to modify the process itself (Fig. 1B) [1, 2].

**Figure 1:**
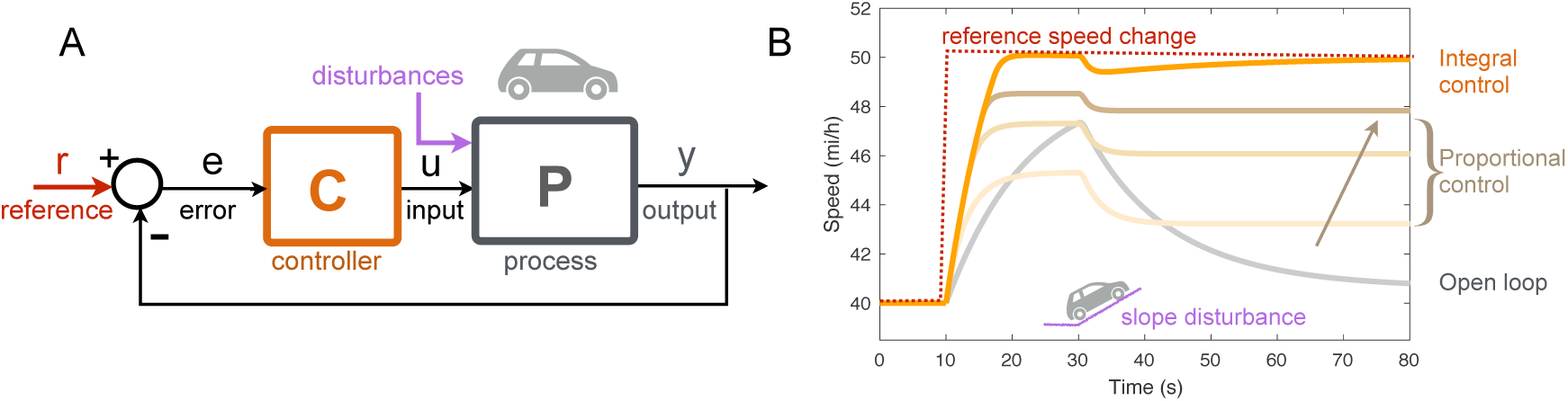
Feedback control enables reference tracking, adaptation, and design of the kinetic response. A: Block diagram of a car cruise control system, in which the actual velocity (output y) is compared to the reference velocity and their difference is minimized as the controller computes the control variable u. B: Example simulations of a car cruise control system. A high gain proportional controller, which computes the control variable u as a linear function of the error, operates well but cannot match an integral controller as far as reference tracking and disturbance rejection. (MATLAB Simulink model downloadable online [1].)

A similar approach is desirable when engineering biological processes for homeostasis, *i.e.* the ability to maintain a certain property despite environmental disturbances and unknown reaction rates. For instance, the activity of a certain enzyme in a large metabolic network may have to be kept constant to maximize flux despite variable environmental conditions [3]. A molecular control system could also be used to tune (and maintain) the expression kinetics and the average level of a protein of interest depending on external stimuli, without having to tune the strength of its promoter. For example, the expression level of a receptor or pump may have to rapidly adapt to the concentration of their target molecules [4].

Feedback control of gene expression has been successfully implemented to achieve both classes of behaviors (homeostasis and response design), for applications that range from regulation of cell density to biofuel production [5, 6, 7]. Most genetic feedback control systems are built by introducing a negative autoregulation loop via a suitable transcriptional repressor. Negative autoregulation yields a stable steady-state that can be prescribed by designing the repression threshold and cooperativity, and it can be seen as a biomolecular controller. Unfortunately, this type of controller requires very careful tuning, and its performance deteriorates if the parameters of the process are uncertain, like proportional controllers in engineering applications (Fig. 1B); further, the controller must be redesigned to change the, steady-state or the kinetics of the closed-loop system (for instance, by varying promoter strength tra nscription rate, or half-life of the repressor).

These challenges appear to be absent from many (generally complex) mechanisms that control natural pathways. Processes such as as chemotaxis, the osmotic response, and even many electrophysiological responses are known to be extremely robust and maintain their set-point rejecting disturbances [8, 9, 10]. Mathematical modeling has revealed that this ability to maintain a set-point is due to the presence of an integrator in the system. Similarly, in traditional engineering fields an integral controller guarantees that the process output eventually matches perfectly the desired reference (zero steady-state error) and that disturbances are rejected, a performance that cannot be achieved with proportional controllers (Fig. 1B). Recent theoretical work has explored the use of molecular sequestration to build molecular controllers that perform integral action [11]. Ongoing experimental research aims at building sequestration-based integral controllers using sigma and anti-sigma factors in *E. coli* [12]; this research builds on ample evidence that sequestration is suited to achieve concentration reference tracking, with implementations that go beyond sigma and anti-sigma factors [13, 14] and range from nucleic acid networks *in vitro* [15], to protein [16] and RNA-based sequestration [17, 18]. A significant challenge in implementing integral action with sequestration alone is posed by the presence of dilution, which introduces steady-state error (leaky integration) [19]. Because a key feature of molecular sequestration is that it can yield an ultrasensitive response [20], we were prompted to examine the role of ultrasensitivity in the design of molecular controllers (not necessarily relying on sequestration) that can achieve integral action.

Here we identify general design principles to build well-defined molecular controllers that a) guarantee tracking of a tunable reference or set-point, and b) achieve integral or quasi-integral action, namely the set-point is tracked with zero or asymptotically zero error. We show that this is possible as long as the controller steady-state input-output map has the following general properties: (1) ultrasensitivity, (2) tunable response threshold, and (3) low sensitivity to variations in the parameters and to the presence of downstream components. We introduce a simple reaction network, which we name Brink motif, that enjoys these three properties; the motif combines molecular sequestration and an activation-deactivation cycle, and presents a response akin to zero-order ultrasensitivity (without operating in a saturated regime) [21]. We provide numerical application examples where the Brink controller is used to regulate an *in vitro* network, a gene expression process, and a biofuel production system.

## 2 Methods

Throughout the manuscript, we indicate chemical species with capital letters (*e.g. A*) and their concention with the corresponding lowercase letters (*e.g. a*).

### 2.1 Design principles for a quasi-integral molecular controller

The general architecture of a closed-loop biomolecular feedback system is shown in Fig. 2A. Two subsystems, a biomolecular controller (*C*) and a biomolecular process (*P*), are interconnected via species *U* and *Y*, forming a negative feedback loop. In the presence of an integral controller, the concentration of the output of the biomolecular process *Y* should be identical to the concentration of the reference species *R*. We propose a simple architecture to achieve quasi-integral performance at steady-state, illustrated in Fig. 2B: first, we require that the input-output steady-state maps of the controller (orange line) and of the process (black line) intersect at a single point, which is the only admissible steady-state of the closed-loop system. Second, we require that the controller input-output map be ultrasensitive, *i.e.* the controller steady-state output concentration should increase steeply when its input is larger than a certain threshold. If the controller output response is ultrasensitive, and its threshold is set by the reference species concentration *r*, then the steady-state of the closed-loop system must fall in a neighborhood of *r*: the more ultrasensitive the controller response, the closer the steady-state *y* is to the reference *r*. Even if the process input-output response is uncertain or affected by perturbations (gray area), the closed-loop equilibrium is guaranteed to be close to the desired reference, as long as the controller is not operating in saturated regime (*u* ≪ *r* or *u* ≫ *r*). This simple architecture should yield a robust closed-loop system that a) tracks changes in the reference input *r* (which determines the controller threshold), and b) handles uncertainty and rejects perturbations on the process parameters. Ultrasensitivity guarantees that the closed-loop equilibrium approaches the reference in the presence of perturbations: by increasing the steepness of the controller map we can decrease the steady-state error. Zero steady-state error could be achieved in the limit, if the controller map is a step function, and the controller can perform as an integral controller without including explicitly a reaction that performs integration. However, we need to discuss one important requirement: that the steady-state be stable, *i.e.* that the system’s output *y* converges to *r*, and returns to *r* in the presence of temporal perturbations (disturbances).

**Figure 2:**
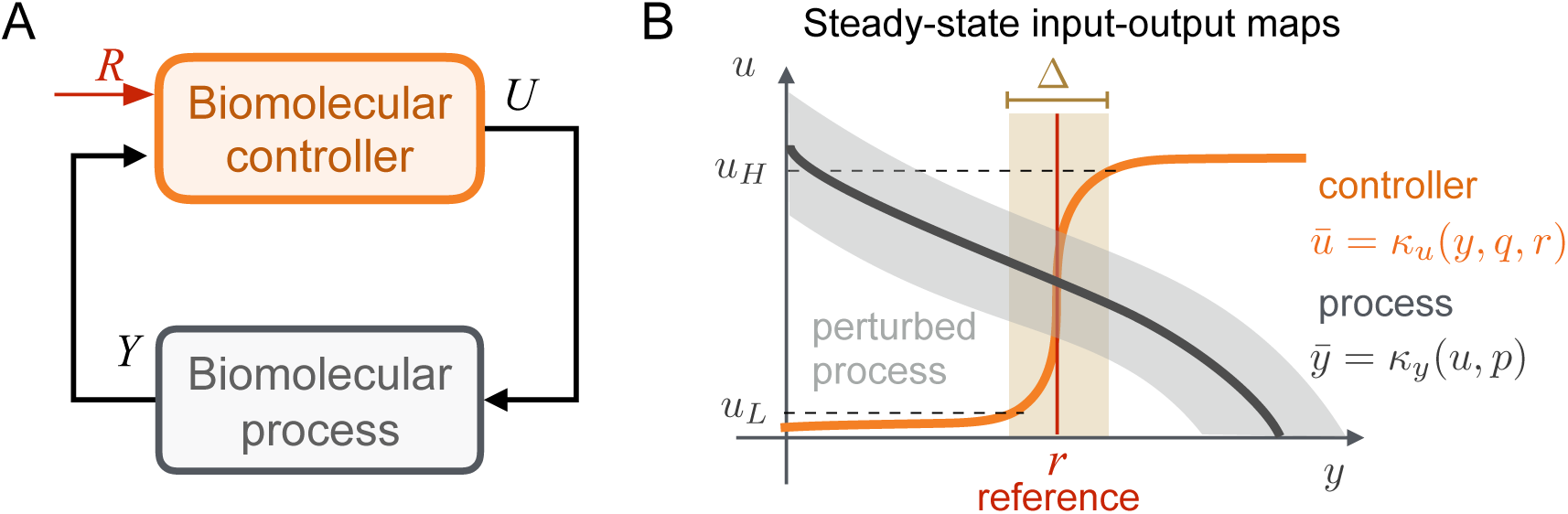
An ultrasensitive controller ensures robust closed-loop performance. A) General architecture of a closed-loop molecular system including the target system to be controlled and the controller module. B) The output equilibrium of the closed-loop system is determined by the intersection of the steady-state maps of the controller (orange) 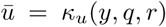, and of the process (dark gray), 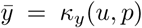; q and p represent parameters characterizing respectively the controller and process. Ultrasensitivity of the controller guarantees that the equilibrium falls in a neighborhood of the threshold or reference (r), characterized by an error Δ(*u, q*) that depends on the controller parameters q (orange shaded area). Reference tracking is achieved even when the process map is uncertain or subject to perturbations in the parameter vector p (gray shaded area).

#### 2.1.1 Assessing closed-loop stability

We consider single-input, single-output molecular processes interconnected to a controller within a feedback loop, like the architecture in Fig. 2A:

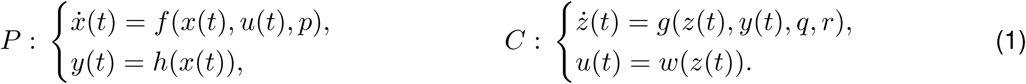

Here, *x* and *z* are concentration vectors, and *y* and *u* are species that interconnect process and controller. The parameters characterizing each system are denoted as vectors *p* and *q*; *r* is a scalar reference (external input to the controller). We will assume that process and controller both admit a unique stable steady-state, which can be found by solving equations *f* (*x, u, p*) = 0 and *g*(*z, y, q, r*) = 0 (for fixed values of parameters and inputs). From these steady-state equations, one can derive steady-state input-output maps:

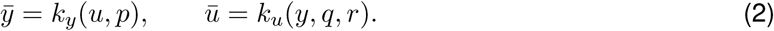

The steady-state values 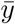 and 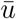 are determined by the intersections of these two maps. Having a unique equilibrium is beneficial in a closed-loop system (since the presence of multiple equilibria would make it more challenging to reach the desired one). A simple way to guarantee the existence of a unique equilibrium is to ensure the input-output maps are monotonic functions: one of them must be increasing, the other decreasing (Fig. 2B); in other words, we require that 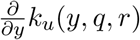 and 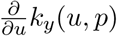 have opposite signs (for example, 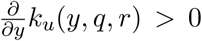 and 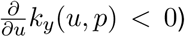). This guarantees that the overall feedback loop is negative.

Given a unique equilibrium, there are many ways to identify its stability. The most direct approach is to linearize the closed-loop system (1) and examine the eigenvalues of its Jacobian matrix evaluated at the equilibrium point [22]. Analytical conditions for stability may be found in models with few variables, but this approach is impractical for realistic, large systems, whose stability must be assessed numerically.

The theory of monotone systems by Angeli and Sontag offers useful tools in this context [23, 24] because many biological models are monotone systems in a broad range of operating conditions, a property that can be verified by qualitatively inspecting their Jacobian matrix [23]. Closed-loop stability can be tested analytically with a loop gain condition depending on the complexity of the models (if they are available), numerically, or even experimentally from steady-state data. However, loop gain conditions are only sufficient, and they are bound to fail in a situation where the controller map is high gain. Yet, monotonicity of the process and controller is useful to guarantee existence and monotonicity of input-output maps, as well as to rule out certain classes of dynamic behaviors [25].

In the examples considered here, we assess closed-loop stability relying on a combination of methods that include linearization and monotone systems theory. When possible, we provide bounds on relevant parameters to ensure stability.

#### 2.1.2 Controller requirements for quasi-integral behavior

The equilibrium of the closed-loop system is determined by the intersection of the input-output static maps shown at equations (2). Perturbations of the process parameters (vector *q*) may cause a shift in the input-output map (gray area in Fig. 2B). However, as long as the maps are monotonic, their intersection is still unique and falls near the controller threshold 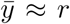 on the horizontal axis, while 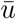 takes any value inside the physically acceptable range of the controller. Therefore, if the controller input-output static map is ultrasensitive like in Fig. 2B, we claim it is possible to write the process steady-state as:

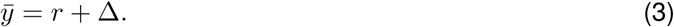

Expression (3) highlights that the difference between the steady-state output *y* and the reference *r* depends on the steepness of the off-on transition of the controller (*i.e.* its ultrasensitivity), which can be quantified with the term Δ (shaded orange area in Fig. 2 B): in the limit of Δ *→* 0 one would achieve perfect integral control. It is reasonable to expect that the deviation term Δ depends largely on the parameters of the controller *q*, rather than the parameters of the process, due to the shape of the ultrasensitive steady-state map of the controller which forces the equilibrium point to be in a neighborhood of the reference. In addition, the equilibrium tracks the reference provided to the controller, and perturbations on the process are rejected. If an expression for Δ is available, it is possible to find lower and upper bounds to the steady-state error 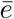, by evaluating function Δ at some chosen threshold saturation values *u*_*L*_ and *u*_*H*_ of the controller: 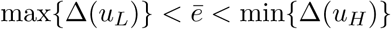.

#### 2.1.3 An ultrasensitive controller operates correctly in its non-saturated regime

We just described how an ultrasensitive molecular controller can help track a reference (which determines the controller threshold) and reject perturbations: correct operation is however guaranteed only if the concentration of controller species *u* does not saturate [26]. If the process map intersects the controller map near saturation, it is not possible for the controller to adjust *u* as required to maintain the reference equilibrium: in this case, the process is not “controllable”^a^. If the controller input-output map 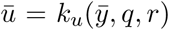 is known, we can formulate a criterion to identify the range of process equilibria 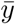 that are controllable:

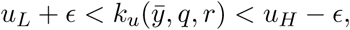

where *u*_*L*_ and *u*_*H*_ are thresholds for the controller. These thresholds should be selected as a perfor<u>mance</u> specification: if *u ≤ u* _*L*_, and *u ≥ u*_*H*_, the controller operates too close to s aturation, and the steady-state deviation Δ from the reference is too large; *E* is a user-defined “safety distance” from the thresholds *u*_*L*_ and *u*_*H*_. Given an input-output map 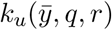 in polynomial form, these inequalities could be used to find bounds on the reaction rate constants that will satisfy the specifications, and to find whether the specifications are realistic in practice for the chosen implementation of the controller. The inequalities could be examined analytically using methods such as the Routh-Hurwitz or Sturm theorem [27, 28] for simple controller networks, or characterized numerically in more complex controllers.

^a^ This is a slight abuse of the traditional meaning of the word “controllability” in control theory.

## 3 Results

Many molecular networks, artificial and native, exhibit an ultrasensitive response (overview in Box 1). However, the threshold and the steepness of the steady-state map of most of these networks are difficult to tune. Because in our approach the threshold of the ultrasensitive controller must be determined by the reference for the process, any implementation should make it possible to vary the threshold over time as a function of an input signal or concentration. We describe a network motif that satisfies this requirement by using molecular sequestration.

### 3.1 The Brink motif produces a tunable, ultrasensitive input-output static map

#### 3.1.1 Brink motif model

A threshold-tunable ultrasensitive molecular network can be built as in Fig. 3A; due to the steepness that is achievable by its static map, we name this network “Brink” motif. The motif has two inputs, an activator species *A* and an inhibitor species *I*, which respectively control the activation and deactivation of downstream species *U*, the output of the motif. While *U* can be in active or inactive state *U* ^*\**^, we assume its total concentration is constant or that it presents negligible fluctuations (*i.e.* the total concentration changes very slowly relative to the other species in the system). The inhibitor *I* produces species *R*_*I*_, which binds to and inhibits *U* by forming the inactive complex *U* ^*\**^. The activator *A* produces species *R*_*A*_, which reactivates *U* ^*\**^ by removing *R*_*I*_ from the complex *Y* ^*\**^, thereby converting *U* ^*\**^ back to *U*. Species *R*_*A*_ and *R*_*I*_ bind to each other (molecular sequestration) to produce a waste complex which does not interfere with the rest of the circuit. In addition, *R*_*A*_ and *R*_*I*_ degrade at a first order rate. The list of reactions is:

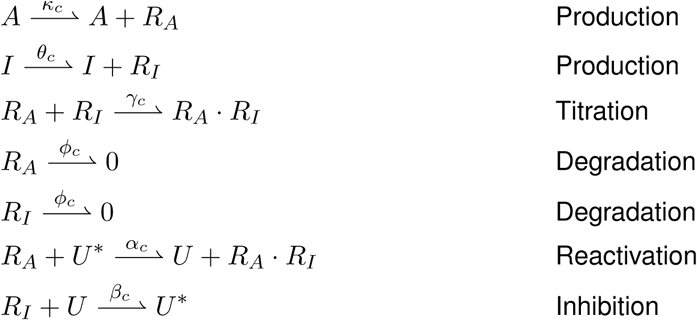

**Figure 3:**
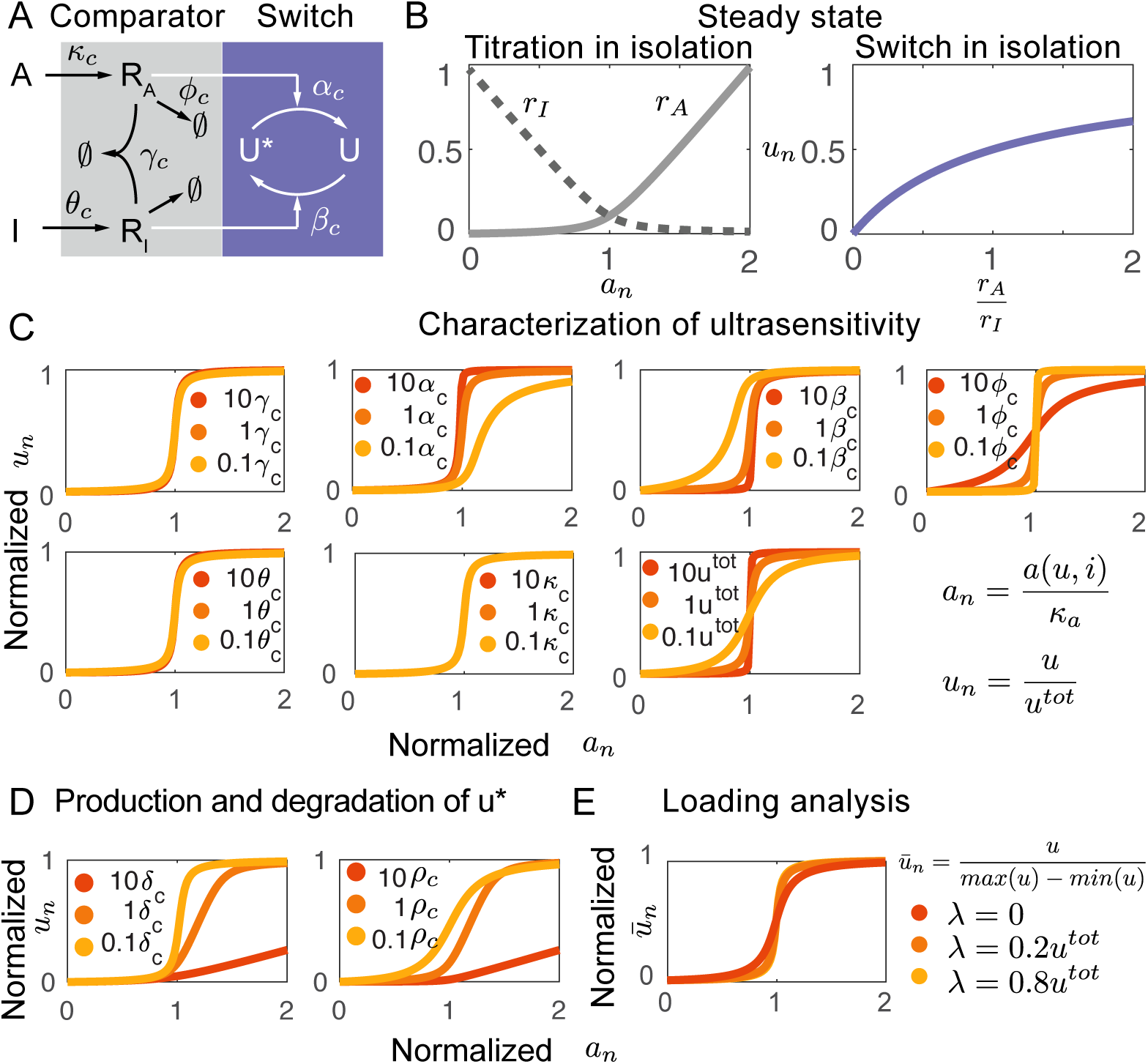
Overview of the Brink Controller motif. All numerical simulations are done using the nominal parameters in Table 1. A) Summary of the reactions defining the Brink Controller motif. The inputs of the motif are species A (activator) and species I (inhibitor); the output of the motif is species U. B) Simulation of the steady-state input-output map of the comparator and switch elements composing the brink motif. Here, the concentration of input I is taken as the reference, while the concentration of input A varies. The concentration of A, a_*n*_, is normalized with respect to the threshold. C) Sensitivity analysis of the input-output static map of the motif. D) Simulation of the input-output map in the presence of production and degradation of the output species; ultrasensitivity is preserved as long as production and degradation rate constants are sufficiently small. The nominal value of δ_*c*_ = 0.77 10^*-*4^ = *ϕ*_*c*_*/*5*, and ρ*_*c*_ = 0.5*δ*_*c*_. E) The Brink motif input-output map, normalized with respect to the output range, is insensitive to the presence of a downstream load.

###### Box 1. Overview of ultrasensitive mechanisms in molecular biology

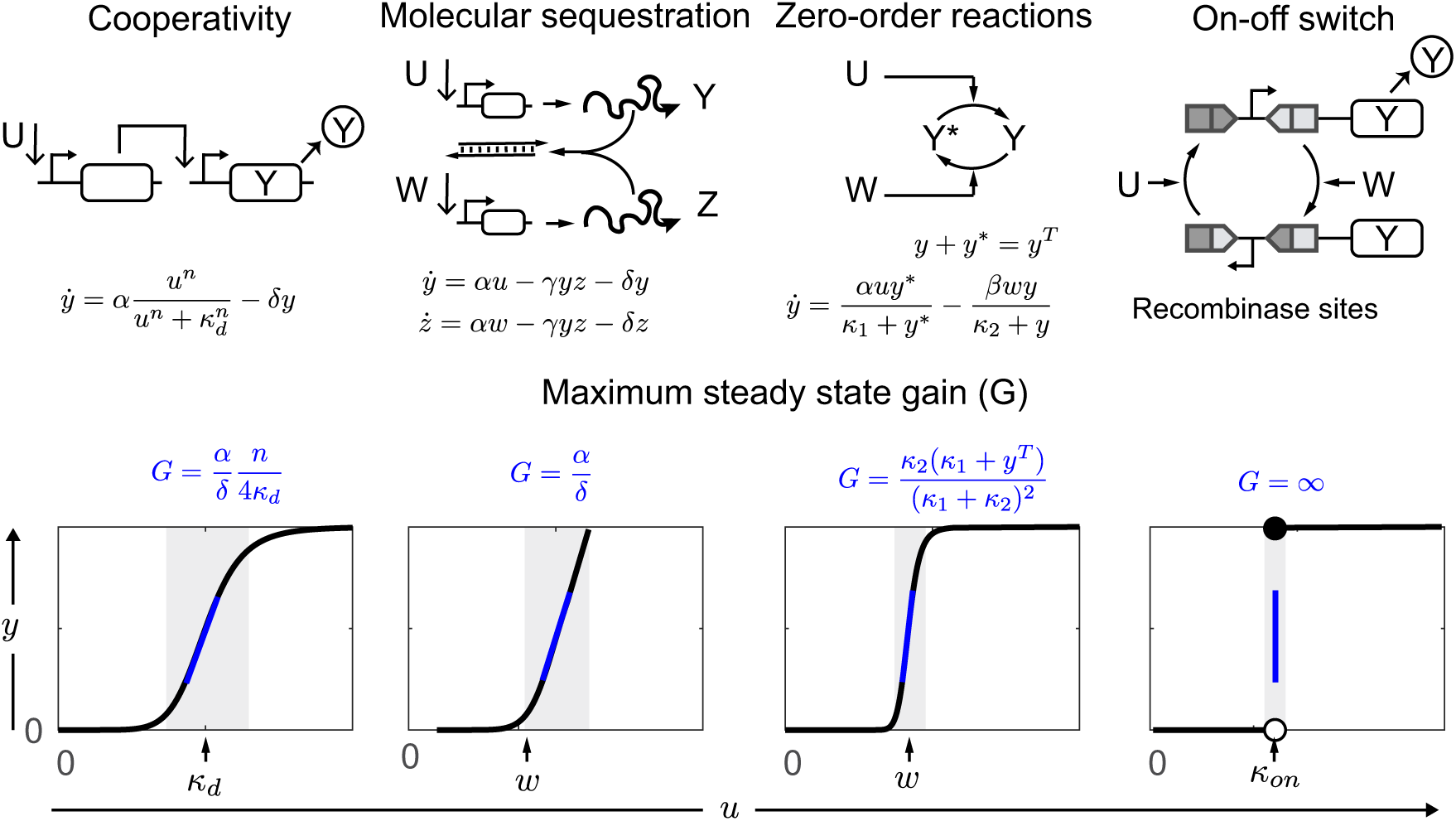

Ultrasensitivity occurs when the steady-state output of a molecular pathway exhibits a sharp increase once the input exceeds a certain threshold. This behavior can be achieved with four classes of mechanisms: (1) Cooperativity; (2) Molecular sequestration; (3) Covalent modification cycles (zeroth-order ultrasensitivity); and (4) On-off switching. Above we describe example ultrasensitive modules relying on these mechanisms via dynamic scalar equations, and we compute their maximum output static gain *G*, *i.e.* the maximum achievable slope of the steady-state input/output response [29].

**Table 1:** *Nominal simulation parameters used for the Brink motif and the controlled process. This table is identical to Table 1 in the main paper and is reported here for the readers convenience.*

| Rate | Description | Value | Other studies |
| --- | --- | --- | --- |
| $\theta_c$ (/s) | Production | $7.5 \cdot 10^{-4}$ | $2.710^{-4} - 1$ [45, 46, 47, 48] |
| $\kappa_c$ (/s) | | $5 \cdot 10^{-4}$ | $2.710^{-4} - 1$ |
| $\gamma_c$ (/M/s) | Titration | $3 \cdot 10^4$ | $10^4 - 10^6$ [49, 50] |
| $\phi_c = \phi_s$ (/s) | Degradation | $3.85 \cdot 10^{-4}$ | $10^{-4} - 10^{-3}$ [51] |
| $\alpha_c$ (/M/s) | Activation | $1.2 \cdot 10^5$ | $10^4 - 10^6$ [52] |
| $\beta_c$ (/M/s) | Deactivation | $1.2 \cdot 10^5$ | $10^4 - 10^6$ |
| $\kappa_s$ (nM) | Dissociation constant | 200 | |
| $\alpha_s$ (/s) | Production | $1 \cdot 10^{-3}$ | $2.710^{-4} - 1$ [45, 46, 47, 48] |
| $\psi_s$ (/s) | | $9.6 \cdot 10^{-4}$ | $2.710^{-4} - 1$ |
| $\phi_s$ (/s) | Degradation | $3.85 \cdot 10^{-5}$ | $10^{-4} - 10^{-3}$ [51] |
| $\delta_s$ (/s) | | $3.85 \cdot 10^{-4}$ | $10^{-4} - 10^{-3}$ |
| $n_s$ | Cooperativity | 2 | $1 - 5$ [53, 54, 55, 56] |

1. Cooperativity is the result of binding-dissociation reactions of multiple molecules at equilibrium, and is quantified by the cooperativity (Hill) coefficient *n* [30, 29]. In a process regulated cooperatively by its input (see equations above), the output static gain *G* is proportional to the output production rate *α* and the input cooperativity coefficient *n*, and it is inversely proportional to the degradation rate *δ* and the apparent dissociation rate *κ*_*d*_. The gain *G* could be increased by cascading cooperative modules (which yield a higher Hill coefficient *n*), which however increases the system complexity and worsens noise propagation [31]. The response threshold of cooperative modules *κ*_*d*_ is also difficult to tune, because molecular association-dissociation rates are not easily modified.

2. Molecular sequestration can generate an ultrasensitive response via stoichiometric interactions [20, 32, 33]. In the example ultrasensitive module above, the gain *G* is proportional to the production rate *α* and inversely proportional to the degradation rate *δ* of the components (SI Section 2.2). The input *w* determines the threshold of ultrasensitivity, which is therefore easily tuned. If the degradation *δ* becomes negligible, then the system will produce an infinite gain *G*, yielding perfect integral action as proposed in the antithetic feedback controllers [11]; in some cases, time-scale separation arguments can allow neglecting *δ* [19]. However, experimental characterization of molecular sequestration processes *in vivo* and *in vitro* suggest that it is difficult to achieve a high static gain on a non-logarithmic scale [34, 35].

3. Zero-order ultrasensitivity occurs when enzymes operate in a regime of saturation. creates a higher gain than molecular sequestration, and it requires saturation of enzymes. The static gain of this example module has been computed in [36, 37]. The input *w* can be used to easily tune the output response threshold. A push-pull motif of a covalent modification cycle was reported to achieve zero-order ultrasensitivity *in vitro* [38], although this behavior is hard to find in nature [36]. This mechanism satisfies our two of the requirements for quasi-integral action (ultrasensitivity and tunable threshold). However, ultrasensitivity is compromised when the circuit is connected to a downstream process [37].

4. Systems with a nearly digital on-off behavior achieve a nearly infinite static gain, as there is a sharp transition between two steady-states. This behavior can be achieved by bistable switches, however the switching threshold usually depends on a combination of the network parameters and may be difficult to tune [29]. Recently, a recombinase protein switch was used to achieve a sharp output transition with nearly infinite gain [39, 40, 41]; in this system the threshold is however difficult to tune as it depends on the dissociation constant of protein-DNA. In addition to being difficult to tune, thresholds may also be asymmetric and depend on the switching direction (hysteresis).

Using the law of mass action, from these reactions we obtain an ordinary differential equation (ODE) model for the Brink motif:

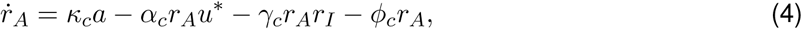

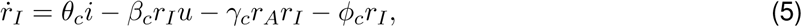

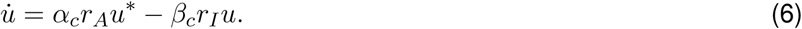

Within the Brink motif we can identify two subsystems: a comparator and a switch (Fig. 3A). The comparator relies on molecular sequestration (or titration), whose purpose is to ensure that only the most abundant species between *I* and *A* has a prevalent regulatory effect on *U* (Fig. 3B). The switch reactions shifts the balance between *U* and *U* ^*\**^, depending on the outputs of the comparator subsystem (Fig. 3B). To achieve the desired operation, one of the inputs (*I* or *A*) should be kept constant, while the other is allowed to vary over time. When used as a controller, the species maintaining constant concentration operates as the reference signal, while the time-varying species is the signal to be controlled.

#### 3.1.2 Ultrasensitivity conditions at steady-state

We derive expressions for the input-output static map of the Brink motif, and we obtain analytical conditions to guarantee ultrasensitivity of the map.

First, we consider the case in which *I* is kept constant and acts as a reference input to the module, while *A* can vary. We derive equilibrium conditions by setting equations (4)-(6) equal to zero. Finding the controller output *u* as a function of *a* and *i* is a long and tedious procedure. However, we can easily express *a* as a function of *i* and *u* in closed form:

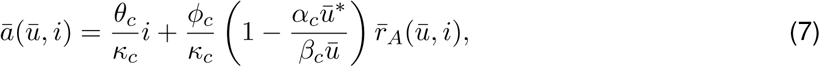

Where

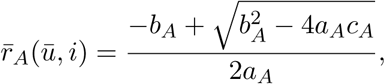

and 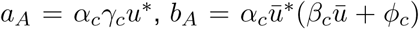 and 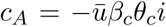. Equation (7) describes the input-output steady-state map of the model (4)-(6) and can be rewritten in a form similar to Equation (3):

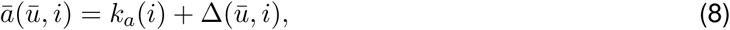

with 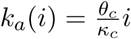 and 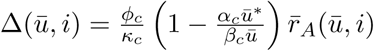. Expression (8) is useful because it allows us to clearly break down 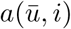 as the sum of a threshold *k*_*a*_(*i*) depending on input *i*, and of an additional term 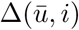.

Are there parametric conditions that make the input-output static map (*u* vs *a*) ultrasensitive? We can answer this question by noting that If the motif is ultrasensitive, small changes in *a* correspond to large changes in *u* when 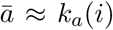. Therefore we focus on term 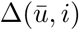, and we look for parameter combinations that make 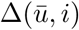 small, for arbitrary values of *u ∈* [0*, u*^*tot*^]. Within term 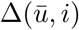, the factor 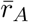 (defined earlier) is small if 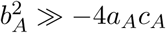, *i.e.* if:

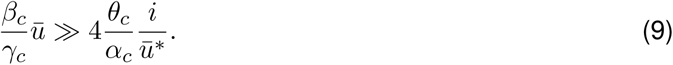

If we assume that the sequestration reaction is fast, thus *γ*_*c*_ is large, then condition (9) is satisfied if the switching parameters *α*_*c*_ and *β*_*c*_ are also large. In addition, the term 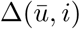 is proportional to the ratio between degradation and production rates of *r*_*a*_, *ϕ*_*c*_*/κ*_*c*_: a small degradation rate constant *κ*_*c*_ promotes ultrasensitive behavior. In the Supplementary Information file (SI) we derive similar ultrasensitivity conditions in the case in which the production rates of *R*_*A*_ and *R*_*I*_ are nonlinear Hill-type rates that exhibit saturation, which may be more realistic in practice (SI Section 3.5).

In summary, we achieve ultrasensitivity when the sequestration and switching rates are sufficiently fast. In addition, the threshold of the ultrasensitive response 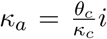 can be tuned linearly with the input *i*. A slow degradation rate *ϕ*_*c*_ also promotes utrasensitivity.

Our derivations are supported by the numerical evaluation of expression (7) in Fig. 3C, which was done using the parameters reported in Table 1. The *x* axis of this plot uses a threshold-normalized input 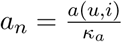. The nominal values of titration rate *γ*_*c*_ and switching rates *α*_*c*_ and *β*_*c*_ are chosen to be in a realistic range for nucleic acid systems, and are on the same order of magnitude. Ultrasensitivity is not drastically affected if we change *γ*_*c*_ within one order of magnitude of the nominal value; however, variations of *α*_*c*_ and *β*_*c*_ in the same range make the transition less sharp. As predicted by our analytical approximation, a small degradation rate *ϕ*_*c*_ improves ultrasensitivity. Changes in *θ*_*c*_ and *κ*_*c*_ primarily affect the threshold *κ*_*a*_, however because our plot uses a threshold-normalized input these effects are not visible.

In SI Section 3.3 we follow similar steps to find the input-output mapping when the inhibitor *i* is varied, while the activator *a* is constant and determines the threshold for the input-output map. Consistently with the analysis above, we find that large *γ*_*c*_, *β*_*c*_ and *α*_*c*_ promote an ultrasensitive response. The threshold *k*_*i*_ can be tuned linearly by the concentration of *a*, similarly to the case considered earlier. Numerical analysis (not reported for brevity) shows that the dependence of ultrasensitivity on parameter variations is analogous to what shown in Fig. 3C.

The expression of the input-output steady-state map of the Brink motif model can be compared to a Hill function having the same threshold maximum slope; without loss of generality, we consider the motif operating as an activated network (activated by input *a*, while species *i* is held constant). The output map of the motif is equivalent to a Hill function: 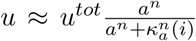, with 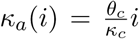, and 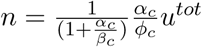. Details on the derivation of these expressions are in SI Section 3.5. When operating in a quasi steady-state regime, the model of the Brink motif could be reduced to a single (static) equation in which *u* is determined by the above Hill function.

#### 3.1.3 The Brink motif is an unconditionally stable, monotone system

The model of the Brink motif is a monotone system and is unconditionally stable, properties that are verified in Propositions 2–4 in SI Section 3. These features can be verified in either operation of the system (as an activator or an inhibitor): input-to-state monotonicity follows due to the fact that the Jacobian is a sign-definite, Metzler matrix; stability follows from the fact that the Jacobian is also an irreducible and diagonally dominant matrix. Monotonicity of the input-output static maps is demonstrated using the implicit function theorem, which also allows us to find analytical expressions for the gains (slope) of the maps.

#### 3.1.4 The sequestration module enables error computation and the switching module increases the gain

The first stage of the Brink motif includes a molecular sequestration reaction. Taken in isolation, this reaction operates as a comparator, and at steady-state it computes the difference between its inputs. We consider the case in which *a* is a time-varying input, and *i* is held constant (the same reasoning can be followed for the case in which *i* varies and *a* is constant). At steady-state, given a constant inhibitor concentration *i* and a fast sequestration rate, the output of the comparator can be approximated as:

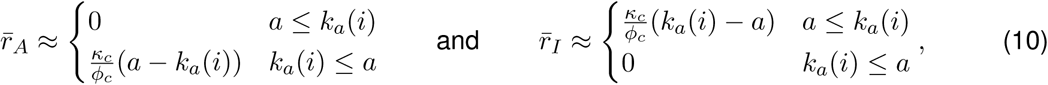

with a threshold 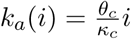. The complete derivations are in SI Section 2. The threshold is proportional to the input *i*, *i.e.* the threshold is tunable by setting the concentration of inhibitor. For special cases where *θ*_*c*_ = *κ*_*c*_, the threshold is equal to the input *i*. This input-output map is plotted in Fig. 3B.

When *a* is larger than the threshold *k*_*a*_(*i*), the steady-state output of the comparator *r*_*A*_ is proportional to *a - k*_*a*_(*i*), which can be interpreted as the error between input *a* and the scaled input *i* with a gain 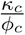. When *a* is smaller than the threshold, the steady-state output *r*_*A*_ is approximately zero. In contrast, the steady-state value of *r*_*I*_ is proportional to *k*_*a*_(*i*) - *a* when *a* is smaller than the threshold, and it is almost zero when *a* is larger than the threshold. The role of the sequestration reaction is thus to create a first layer of input processing that behaves like a diode in electronic circuits, and is characterized by the threshold *k*_*a*_(*i*), so that 1) *r*_*I*_ is almost zero when *r*_*A*_ is being produced, and 2) *r*_*A*_ is almost zero when *r*_*I*_ is being produced. An example simulation is in Fig. 3B, left. A thorough analysis, based on the system’s frequency response, shows that the comparator works well also on time-varying inputs *a*(*t*) and *i*(*t*), as long as they evolve on a timescale slower than the degradation rate *ϕ*_*c*_, as we show in Section 2 of the SI file.

Within the Brink motif, the comparator module is followed by an activation-inhibition cycle in isolation which receives inputs *r*_*A*_ and *r*_*I*_, and produces the output *u* of the comparator. The equilibrium value of *u*, for constant inputs *r*_*A*_ and *r*_*I*_, is given by the following Michaelian function:

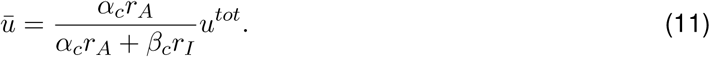

An example simulation showing the input-output map of this module is in Fig. 3B, right.

Based on the expressions we just derived, we can qualitatively explain the overall behavior of the Brink motif when the comparator module and the activation-inhibition cycle are interconnected. When the input *a* is smaller than the threshold *k*_*a*_(*i*), *r*_*A*_ is almost zero and *r*_*I*_ is large, thereby pushing the value of 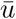 to approach zero according to equation (11). In contrast, when *a* is larger than the threshold *k*_*a*_(*i*), *r*_*A*_ is large and *r*_*I*_ is almost zero, therefore 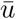 approaches *u*^*tot*^. From expression (11) it is also apparent that if the switching rate constants *α*_*c*_ and *β*_*c*_ are large, the ultrasensitive response becomes sharper.

#### 3.1.5 Slow production and degradation of the output species do not reduce ultrasensitivity

Although their level may be calibrated to stay nearly constant, most regulatory molecules in the cellular environment are dynamically produced and degraded. Thus, we examine the effects of production and degradation reactions of the output species *U* on the behavior of the Brink motif. We consider, without loss of generality, the motif when the inhibitor input *I* is held constant. In this case, we assume that *u*^*\**^ (inactive output) is produced at a rate constant *ρ*_*c*_ and degraded with a rate constant *δ*_*c*_, while the dynamics of *r*_*A*_ and *r*_*I*_ are unchanged:

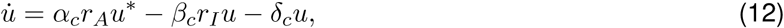

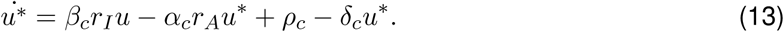

First, we note that *u*^*tot*^(*t*) = *u*(*t*) + *u*^*\**^(*t*), and 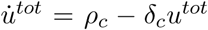, whose equilibrium value is *ρ*_*c*_*/δ*_*c*_; if production and degradation are slow, it would be sensible to simply assume that *u*^*tot*^ *≈ ρ*_*c*_*/δ*_*c*_. Yet, it is useful to derive the input-output map of the Brink motif:

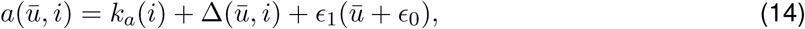

in which *k*_*a*_(*i*) and 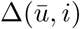 are defined consistently with expression (8), and *∈*_0_ = *ϕ*_*c*_*/β*_*c*_ and *∈*_1_ = *δ*_*c*_*/κ*_*c*_ (complete derivations are in SI Section 3.3). The motif can exhibit an utrasensitive response when 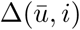 is small; this is achieved when *γ*_*c*_ *≫ ϕ* _*c*_, *α* _*c*_ *≫ δ* _*c*_, and 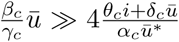, which can be satisfied if the switching rate constants *β*_*c*_ and *α*_*c*_ are large. While these requirements are similar to what derived earlier, we note that introducing production and degradation of the output results in a new term 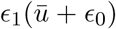 in the input-output map. While *∈* _0_ is negligible as long as *β*_*c*_ is large, the term 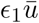 can only be neglected if we assume that *δ*_*c*_ (the degradation rate constant of *u*) is sufficiently small relative to *κ*_*c*_ (the production rate constant of the activator). These analytical derivations do not provide insights into the effects of *ρ*_*c*_; however, with numerical simulations in Fig. 3D, we show that as long as *δ*_*c*_ and *ρ*_*c*_ are sufficiently small the motif preserves its ultrasensitivity property. A complete sensitivity analysis is reported in SI Section 3.3.

#### 3.1.6 Leaky production introduces a bias in the threshold

Inducible promoters in practice always present a basal level of RNA transcription, which results in some level of expression that is independent from the presence or absence of inducer. This is often referred to as a “leak” in the production of the output [42]. In the Brink network, the presence of a basal level of production of species *R*_*A*_ and *R*_*I*_ could be modeled with constant production rates 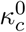 and 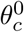:

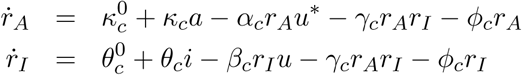

The presence of these terms has an influence on the steady-state map that relates the input and the output of the motif, in particular they introduce a constant bias in the ultrasensitivity threshold (SI Section 3.7):

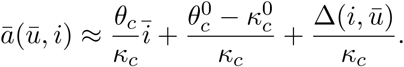

This shift in the threshold can increase the steady-state error, although the transition of the controller curve is still determined primarily by the constant input 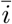 This threshold shift could be reduced by a large transcription rate *κ*_*c*_.

#### 3.1.7 The Brink motif preserves ultrasensitivity in the presence of a downstream load

If the Brink motif is used as a controller in a closed-loop system, its output *u* is expected to bind to at least one downstream component. Thus, a fraction of *u* may be depleted by downstream reactions, which can be considered a load on the controller: in many cases, this is known to cause undesired perturbations on the upstream reactions through a phenomenon called retroactivity [43]. The input-output equilibrium conditions of the system change in the presence of a load, a phenomenon that can result in deterioration of performance [44]. However, the ability of the Brink motif to achieve quasi-integral performance is determined exclusively by the ultrasensitivity of its input-output map, which is not affected by the presence of a load. This can be shown with a model example, in which *u* is depleted by binding to a downstream promoter *g* to produce a species *p* according to these reactions:

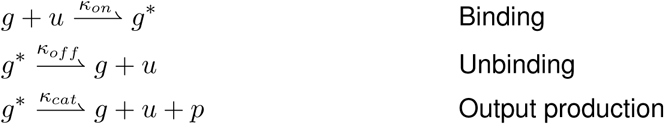

The addition of these reactions to the Brink motif model requires that the mass balance of *u* be updated as:

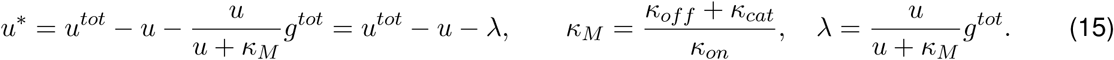

Term *λ*, which captures the loading effects at steady-state, is bounded and has a maximum *λ* = *g*^*tot*^; this term also decreases when *κ*_*M*_ is large relative to *u*^*tot*^, which is occurs when *κ*_*on*_ (the binding rate of *u* to the load) is small.

Equation (15) clearly suggests that the load depletes molecules of *u* thereby reducing the amount of available *u* to downstream processes: as one should expect, simulations show that the presence of the load reduces the range of the input-output map (y-axis), highlighting the limited capacity of the controller to direct downstream processes (SI Section 3.6). However, upon normalizing relative to the maximum range of available *u*, we find that ultrasensitivity of the map is not affected, as shown in Fig. 3E. In other words, the controller is likely to perform well in terms of tracking and disturbance rejection, but there is a limit on the amount of load it can handle. If the concentration of load is low, the loading effect is negligible even in the present of a strong binding site for *u*.

### 3.2 The Brink motif as a closed-loop quasi integral controller

We now consider the Brink motif interconnected with a target process in a closed-loop system. Here we assume that the closed-loop system is stable (verification of this property should be done on a case by case basis). To evaluate its performance, we focus on two criteria: the capacity of the closed-loop system to track changes in reference with small steady-state error, and to reject perturbations of the process parameters. In the rest of this section, we assume the process has a monotonically increasing input-output static map. Thus, we operate the Brink motif as a controller with the activator input as reference *R*, and the inhibitor input is interconnected to the process output *Y* and used as an inhibitor for the motif; this will result in the controller having an input-output map that is monotonically decreasing, guaranteeing that the feedback loop is negative. The output of the controller is *U*, which becomes the input for the process.

The dynamics of the interconnected controller and process are:

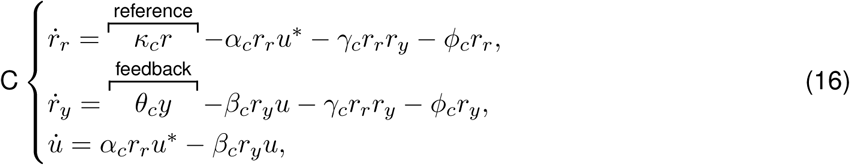

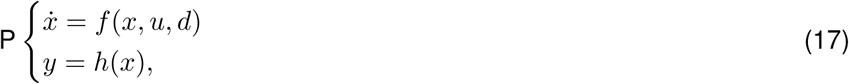

with mass conservation *u*^*tot*^ = *u*^*\**^ + *u*. As derived in equation (8), the output of the process at steadystate is:

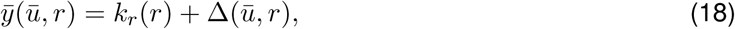

where 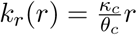 is a scaled reference and 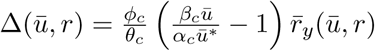.

#### The Brink motif operates as an integral controller near steady-state

When the degradation rate constant *ϕ*_*c*_ is zero, we have 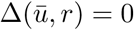, and thus:

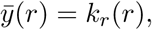

*i.e.* the system exhibits perfect integral action. In some *in vitro* systems degradation can be eliminated or reduced, thus it may be reasonable to assume *ϕ*_*c*_ *≈*0. In contrast, *in vivo* systems always present degradation and dilution (due to cell growth and division), thus *ϕ*_*c*_ *>* 0 and 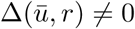. However, sharp ultrasensitivity can yield 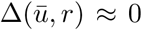 when 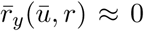, which occurs for large values of *α*_*c*_ and *β*_*c*_, as shown in expression (7). By achieving quasi-integral action, the controller confers to the system the capacity to perfectly track a time-varying reference and to reject disturbances.

Near equilibrium, the Brink motif operates as an integral controller in closed-loop. In SI Section 3.4, we show that if the parameters of the Brink motif satisfy the ultrasensitivity condition (9), then the input-output transfer function of the linearized system (near equilibrium) includes a pole at the origin in the Laplace domain. In turn, this means that the corresponding closed-loop system includes a zero at the origin and is therefore insensitive to step inputs. In other words, the motif as a closed-loop controller rejects small perturbations near equilibrium by performing integral action.

#### The steady-state error of the closed-loop system is bounded

An analytical approximation for the steady-state error can be derived from equation (18):

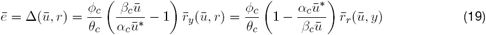

Assuming *α*_*c*_ = *β*_*c*_, we find upper and lower bounds for the error when the equilibrium 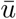 in the interval from 0.1*u*^*tot*^ to 0.9*u*^*tot*^:

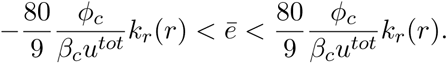

These bounds can be derived from expression (19), recalling that 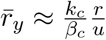 (SI Section 3) and 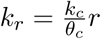.

#### Comparing the Brink motif with a controller based exclusively on molecular sequestration

We have just shown that the Brink motif achieves quasi-integral operation at steady-state when used as a controller in closed-loop. Yet, recent work has shown that sequestration alone can provide integral action, in particular in the absence of degradation or under high-gain conditions [11, 19]. With simulations that use consistent reaction rate constants, we compare the performance of a controller based exclusively on molecular sequestration with the Brink motif (Fig. 4A, yellow and light blue boxes). As an example model process we consider expression of a target protein *y* (Fig. 4A, gray box):

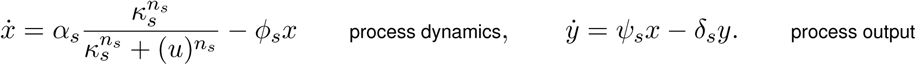

**Figure 4:**
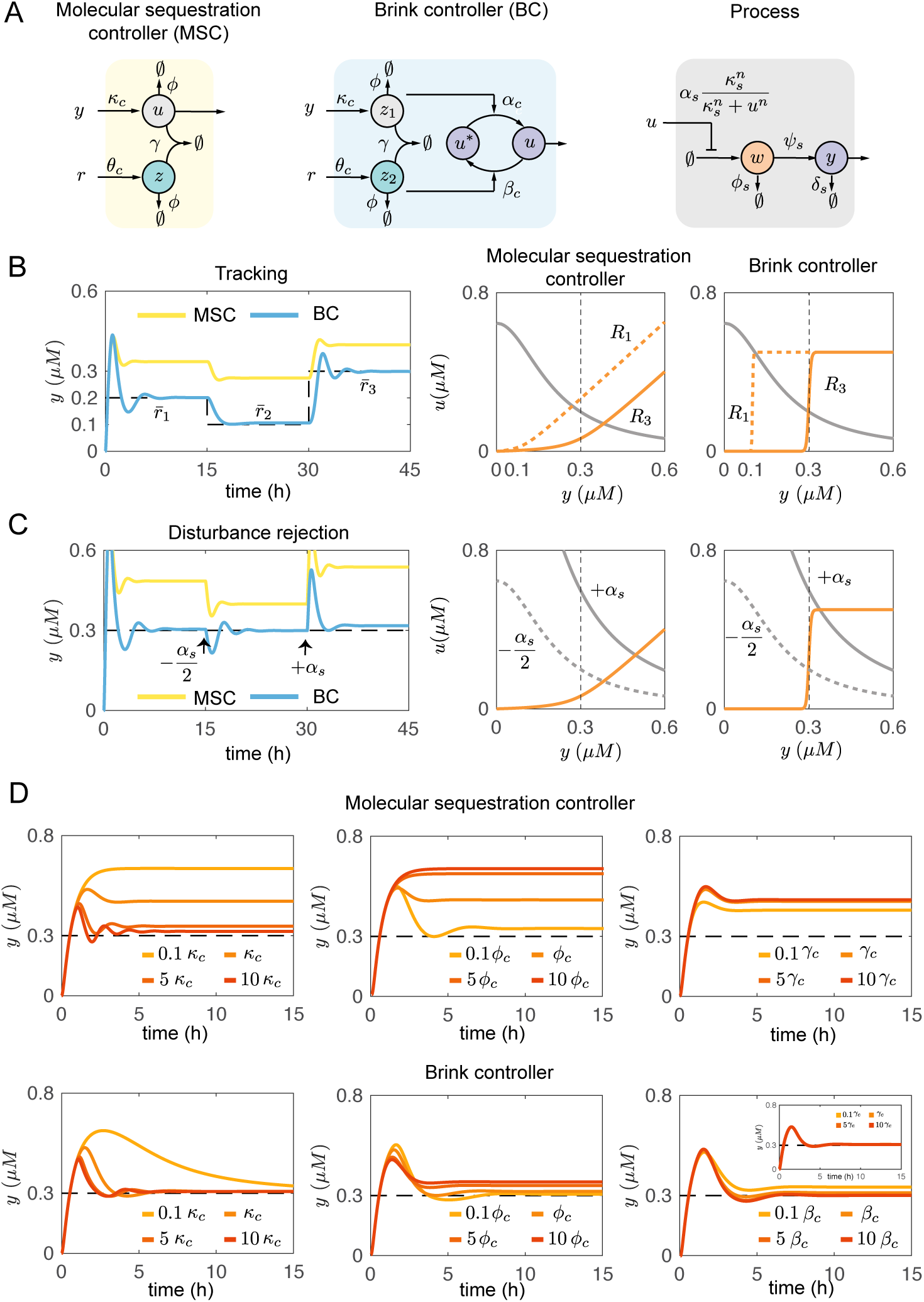
Comparing the Brink controller (BC) with a molecular sequestration controller (MSC) for closedloop feedback. ODEs were integrated using the nominal parameters in Table 1. A: Schematics summarizing the reactions occurring in molecular sequestration controller, in the Brink motif controller, and in the model gene expression process. B: Left: Comparison of reference tracking performance in closed-loop. Right: The equilibrium conditions of controller (orange) and process (gray) confirm that tracking is achieved only in the presence of the Brink network. The ultrasensitivity threshold changes as a function of the reference. C: Left: Comparison of disturbance rejection performance. Right: equilibrium conditions. The process steady-state map changes with variations in the process parameter (α_*s*_ is taken as an example), however in the presence of the Brink network the equilibrium point remains near the reference value. D: Temporal behavior of the process as a single parameter is varied (with respect to the nominal values in Table 1) in each plot. The Brink controller outperforms the molecular sequestration controller in all cases in terms of steady-state error. The closed-loop system is not sensitive to changes in γ_*c*_ when the Brink controller is used (inset in the bottom right plot; all traces overlap); the system is somewhat sensitive to changes in the switching rates α_*c*_ = *β*_*c*_, but the steady state error is small in all cases considered.

The ODEs of each controller are reported in SI Section 3.9; the ODEs are integrated using the nominal parameters listed in Table 1. In Fig. 4B, we report numerical simulations showing that, for the nominal parameters we selected, the Brink motif yields zero steady-state error in response to changes in reference, while sequestration alone does not manage to achieve the goal; we show the solutions to the ODEs (left) as well as the equilibrium conditions (right), which highlight how shifting the reference results in a shift of the controller ultrasensitivity threshold. In Fig. 4C we compare the ability of these controllers to reject disturbances in the parameters. We vary the transcription rate *α*_*s*_, which results in a change of the process equilibrium conditions (right, gray line). The Brink motif maintains the desired reference perfectly rejecting this disturbance, while the sequestration-only controller results in a steady-state offset. In Fig. 4D we study the behavior of the output of the closed loop system when individual controller parameters are varied. To achieve a small steady state error, the molecular sequestration controller requires very large production and sequestration rate constants, and very low degradation rate constant [11, 19]. In contrast, the Brink controller achieves similar performance within a broader range of the same parameters; notably, the Brink-controlled system is insensitive to changes in the sequestration rate *γ*_*c*_, however it presents some sensitivity to variations in the switching rate constants *α*_*c*_ = *β*_*c*_. These simulations suggest that, assuming the nominal parameters in Table 1, one could compare the performance of the molecular sequestration controller to a proportional controller in traditional engineering fields, while the Brink controller operates similarly to an integral controller.

### 3.3 Application examples

We present two potential implementations of the Brink motif based on RNA, and we examine the ability of the motif to control three different molecular processes of increasing complexity: a synthetic transcriptional network, a gene expression (transcription-translation) process, and a biofuel production process.

#### 3.3.1 Implementing the Brink motif using RNA

As the Brink motif relies on sequestration and switching, its implementation would require the ability to design molecules with the capacity to both sequester each other, also bind to and alter the activity of downstream elements. RNA molecules are ideal for these purposes, because RNA-RNA interactions can be rationally programmed based on their sequence content [57, 58], and a variety of RNA aptamers can be used to control activity of proteins [59]. RNA molecules with multiple domains having distinct functions can be engineered with systematic methods [60], and complex RNA-based reaction networks have been amply demonstrated in *in vivo* [61, 62]. In addition to their programmability, RNA regulatory molecules enjoy other important advantages over their protein-based couterpart: a low metabolic burden (because they do not require translation), portability (synthetic RNA molecules are not host-specific), and fast response (production and degradation rates of RNA are generally faster than proteins).

We can estimate the gain of the comparator stage built with mutually sequestering RNA species. The steady-state gain of the comparator *κ*_*c*_*/ϕ*_*c*_ depends only on RNA production and degradation rates. This gain could be crudely estimated in different types of biological cells using average mRNA transcription and degradation rate constants. The life time of mRNA is roughly 5 minutes in *E. coli*, 20 minutes in budding yeast, and 600 minutes in human cells [63]. The average transcription rate in *E. Coli* is between 50 *-* 100 nucleotides (nt) per second [64, 65, 66], and in eukaryotic HeLa cells it is 30 *-*100 nt/s [67], thus we take an average 60 nt/s *≈* 360 nt/min transcription rate. For a short mRNA species that is 100 nt long, which is typical for a synthetic RNA circuit, we could compute the gain as *G* = *κ*_*c*_*/ϕ*_*c*_ = (txn rate*/*mRNA length)*/*(*ln*(2)*/*life time). With the transcription rate estimates above, we obtain *G ≈* 26 for *E. coli*, *G ≈* 104 for buddying yeast, and *G ≈* 3100 for mammalian cells. However, the actual production rate of mRNA is much slower than the transcription rate, because promoter binding and engagement of RNA polymerase is inefficient. A more realistic estimate of average mRNA production in *E. coli* is of roughly 0.4/min [68]; in human cells, the mRNA production rate is estimated to be around 0.96 *-* 1.92/min [66]. This yields an estimated gain *G ≈* 2.8 for *E. coli* and *G ≈* 6.9 *-* 13.8 for mammalian cells. These gains could be increased further by using high copy number plasmids and strong promoters, and may be up to 5-6 times larger during exponential cell growth [69]. Yet, these estimates suggests that the gain achievable by sequestration alone is not sufficient for quasi-integral control, and that it is highly desirable to increase the gain with an activation/deactivation cycle downstream of sequestration.

We propose two implementations of the Brink controller:

##### 1. Aptamer Brink Controller

The reactions are shown in Fig. 5A, right. This implementation is suited in particular for artificial gene networks *in vivo*, but has the potential to be adapted to work inside cells. The comparator stage is built with two RNA species *R*_*I*_ and *R*_*A*_, which are respectively an aptamer and a complementary anti-aptamer. The aptamers can be produced with synthetic templates (*A*, *I*). The switch stage is built using the RNA aptamer (*R*_*I*_) to bind to and deactivate a viral RNA polymerase [70, 71]; the anti-aptamer *R*_*A*_ displaces the aptamer, reactivating the polymerase. The switch stage has been recently demonstrated *in vitro* [52]. While at the moment only two viral RNA polymerases (SP6, T7) have been regulated using aptamers, it is possible to select aptamers against other polymerases or transcription factors using SELEX [72]; natural aptamers regulating RNA polymerase in *E. coli* have been recently discovered [73].

**Figure 5:**
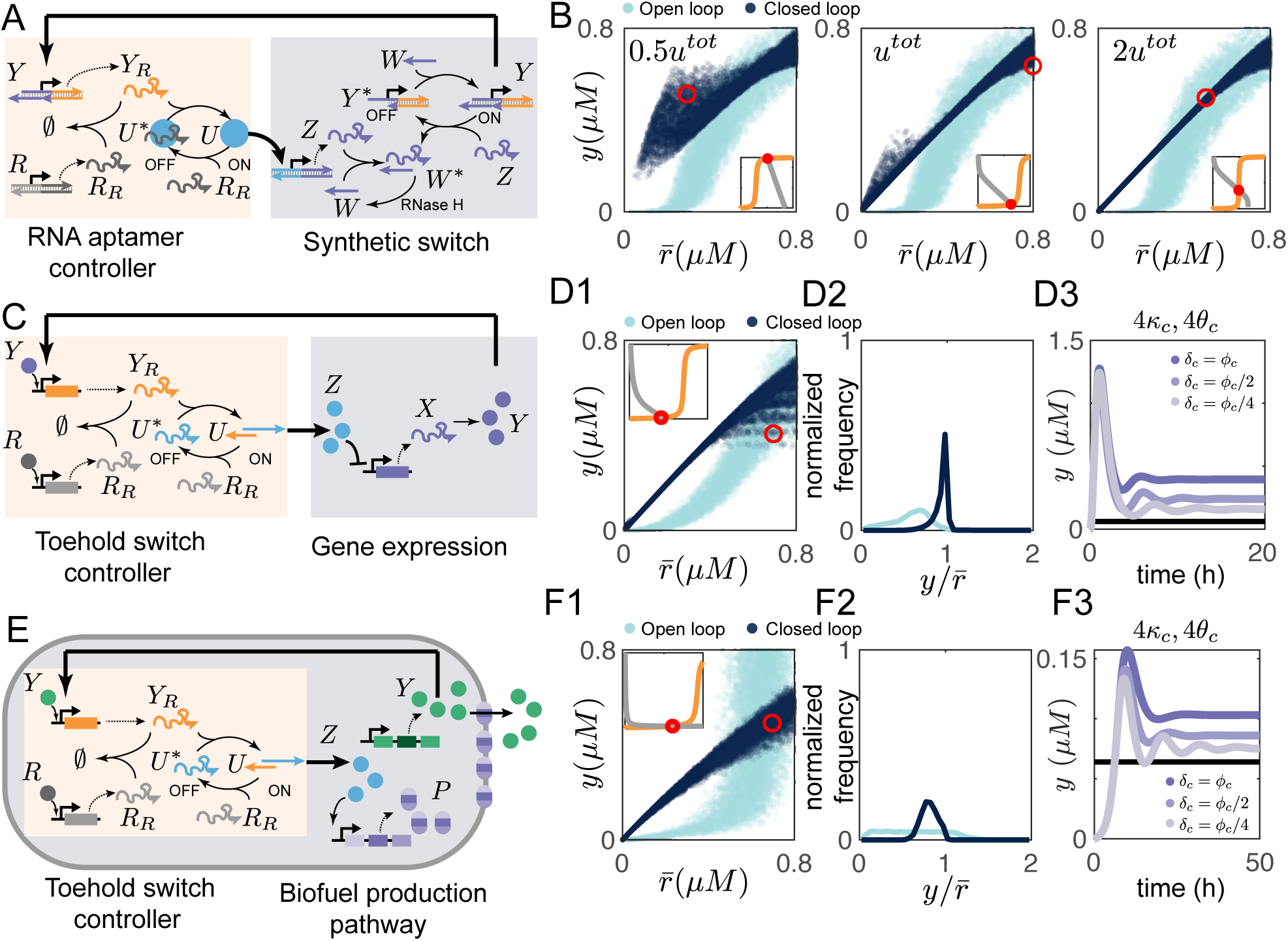
Implementation examples of the Brink controller. A: Schematic of the reactions for an aptamer-based Brink controller applied to the closed-loop control of a synthetic genelet [49]. B: Simulations showing that the Brink controller achieves linear reference tracking (dark blue), outperforming the open loop system (cyan); each circle represents the equilibrium resulting from process parameters randomly perturbed within ± 20% of their nominal values (SI Table S2). The plot includes 10,000 equilibrium points each computed from a random parameter draw. Increasing u^tot^ improves the closed loop system performance. Insets show example equilibrium maps. C: Reactions for toehold-switch Brink controller, used to control gene expression. D1: Steady-state performance of the closed loop (dark blue) and open loop system (cyan); each circle is the equilibrium computed by randomly varying the process parameters (10,000 draws) within ± 20% of their nominal value (SI Table S3). Inset shows an example imput-output map, highlighting that poor performance can occur when the equilibrium falls near the controller saturation regime. D2: Frequency plot showing that most random parameter combinations yield a good tracking performance in the closed loop system. D3: Example simulation showing that in the presence of production and degradation (rate constants ρ_c_, δ_c_) of the controller output species U ^*^, performance of the system can be maintained as long as these rates are slow (here ρ_c_ = 0.5δ_c_) and and the Y_R_ and R_R_ production rates are fast (κ_c_, θ_c_). Reference concentration is in black. E: Toehold switch Brink controller applied to a biofuel production system. F1-F2: We compare the ability of the output of the closed loop and open loop system to achieve a desired reference. The process parameters were randomly drawn in the interval ± 20% their nominal value (10,000 draws); the closed loop system linearly tracks the reference outperforming the open loop system in the majority of parameter combinations (F2). F3: If component U ^*^ is produced and degraded at rates ρ_c_, δ_c_, the controller performance is preserved as long these rates are slow (ρ_c_ = 0.5δ_c_). The reference is shown in black.

##### 2. Toehold Switch Brink Controller

The reactions are shown in Fig. 5B, left. In this implementation, the comparator stage is again built with two complementary RNA species that mutually sequester each other. Species *U* is a toehold RNA switch [61, 74], which is constitutively in its inactive form *U* ^*\**^ because a programmed secondary structure hinders access of the ribosome to the RBS region. Since ribosomes cannot bind to *U* ^*\**^, translation cannot occur. RNA species (*R*_*A*_) is designed to interact with *U* ^*\**^ and convert it to its active form *U*, to which the ribosome can bind and start translation. In our proposed design, *R*_*A*_ should be designed to include a toehold, so that the complementary RNA species *R*_*I*_ could be used to displace *R*_*A*_ bound to *U*, thereby deactivating it. Implementation of this controller should take into account the dynamics of production and degradation of RNA species *U* (or *U* ^*\**^), and we discuss this point in the examples below. Controllers based on toehold switches could be realized for a variety of processes; there are at least 26 orthogonal toehold switches available, with a dynamical over 400-fold, that could be integrated into the genome to regulate up to 12 genes independently [74]; recently these switches were tested in *E. coli* to build logic devices that accept up to 12 distinct inputs [61].

#### 3.3.2 Examples of closed-loop control using the Brink motif

##### Controlling an artificial transcriptional switch

Artificial transcriptional switches have been used to demonstrate a variety of programmable dynamic circuits *in vitro* [49, 44, 75]. Fig. 5A, right, shows the schematic of an *in vitro* synthetic switch (process to be controlled), which receives a single input species *U* and produces an output species *Y*. *Y* is a linear template (genelet) whose promoter is partially incomplete; the promoter region is completed upon hybridization of a single-stranded DNA activator molecule *W*. The template is deactivated by an RNA inhibitor molecule *Z*, which displaces *W* via toehold-mediated branch migration [76]. The input *U* is a viral RNA polymerase transcribing inhibitor *Z* from a distinct (constitutively active) template; *Z* is designed to bind to *Y* and convert it to inactive *Y* ^*\**^ (this occurs by displacement of the activator *W*). In addition, *Z* directly binds to and sequesters *W* converting it to inactive species *W* ^*\**^. We assume that inhibited activator *W* ^*\**^ spontaneously reverts to its active form *W*, due to the presence of enzymes responsible for degradation (*e.g.* RNase H). The total concentrations of *Y* and *W* are assumed to be constant, *y*^*tot*^ = *y* + *y*^*\**^ and *w*^*tot*^ = *w* + *y* + *w*^*\*.*^

The control objective is to maintain a constant active genelet concentration *y*, which should be equal to a specific reference concentration (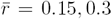 and 0.45*μM*). The circuit operation can be significantly affected by variability in enzyme activity and uncertainty in the value of the reaction rates.

We design an Aptamer Brink Controller to control the concentration of active genelet *y*, as shown in Fig. 5A, left. The controller computes the error (*r - y*): if this difference is positive (insufficient *y*), it reduces the concentration of free active enzyme *u*, thereby decreasing transcription of *z*; if the difference is negative (excess *y*), the controller increases the concentration of free *u* in order to increase the transcription of *z*. The full set of reactions and the corresponding ODE model (controller and transcriptional process) is reported in Section 4.1 of the SI file, where we also show that the closedloop system always presents a unique equilibrium that remains stable in a range of parameter values.

Fig. 5B, top, shows the steady-state behavior of the circuit by comparing the normalized reference value 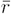 to the output (dark blue circles) relative to the circuit operating in open loop (cyan circles); in these plot, the nominal parameters are randomly drawn in an interval *±* 20% of their nominal value. The controller performs more robustly (with respect to uncertainty in the parameters) when the total concentration of controller output (enzyme *u*^*tot*^) is increased, thereby increasing its range. In the insets, we plot example equilibrium maps corresponding to parameter combinations that yield the desired reference tracking, or fail to perform. These plots confirm that tracking is guaranteed when the input-output maps intersect in the ultrasensitive region of the controller; if the intersection occurs in the saturation region of the controller, reference tracking fails. The nominal parameters used for these simulations are in Table S2. The sensitivity of the closed-loop system to changes of individual parameters is explored with simulations in SI Section 4.1.

##### Control of gene expression

In this case our objective is to control expression of the protein *y* so that it tracks a reference concentration despite uncertainty in the process reaction rates. Fig. 5C, right, illustrates the processes of mRNA transcription and translation of a protein *y* induced by an input transcription factor *z*. Typically, in open-loop the only simple approach to achieve a desired reference concentration *y* is to titrate the inducer concentration *z*.

This system can be controlled using a Toehold Switch Brink Controller, shown in Fig. 5C. The complete model of this closed-loop system is reported in Section 4.2 of the SI file, where we also report all the simulation parameters (Table S3). For this example, we assumed the production rates of *R*_*A*_ and *R*_*I*_ are Hill functions that are more likely to represent the switch behavior; this variation of the model does not affect the controller ultrasensitivity regime (SI Section 3.8). The closed-loop system admits a single equilibrium point, as demonstrated analytically in SI Section 4.2, where we also numerically simulate the effects of variations of individual parameters on the temporal behavior and equilibria of the system.

Simulations in Fig. 5D1 compare the steady-state behavior of the closed loop system (dark blue circles) and open loop system (light blue circle) when the parameters of the process are randomly drawn in an interval that spans *±* 20% their nominal value (Table S3). These simulations show that the closed loop system tracks the reference very precisely in a broad range; for references above *≈* 0.4 *μ*M, some parameter combinations yield a poor tracking performance, as the equilibrium occurs in the lower saturation region of the controller; this is exemplified in the inset plot. However, the controller operates correctly in the majority of parameter combinations, as shown in the frequency plot of Fig. 5D2. The sensitivity of the closed loop system to variations of individual process parameters is numerically explored in SI Section 4.2.

In this controller, *U* and *U* ^*\**^ are RNA species, which must be transcribed and degraded. We model this case introducing a production rate constant *ρ*_*c*_ for *U* ^*\**^, and assume that both *U* and *U* ^*\**^ degrade with a rate constant *δ*_*c*_. To enable correct operation of the controller, we find that *ρ*_*c*_ and *δ*_*c*_ should be sufficiently slow in relation to other parameters, as exemplified by the numerical simulations in Fig. 5D3. Here *ρ*_*c*_ = 0.5 *δ*_*c*_, so that at equilibrium *u*^*tot*^ ≈ 500 *nM*. Consistently with expression (14) derived earlier, to guarantee a small steady-state error we must increase the production rate constants *κ*_*c*_ and *θ*_*c*_ (here assumed to be four times their nominal values in Table S3) of intermediate species *Y*_*R*_ and *R*_*R*_, and ensure a low degradation rate *δ*_*c*_ for the controller output.

##### Microbial Biofuel Density Control

We consider a model of microbial biofuel production system which was proposed in [4]. Biofuel production with genetically engineered microbes promises to expand our renewable energy sources. However, biofuels are generally toxic for microbes and inhibit their growth, thereby limiting the yield of biofuel production. This challenge can be mitigated by introducing biofuel efflux pumps that reduce the intercellular concentration of biofuel and increase the host resilience. Because the over-expression of pumps can negatively affect microbial growth, there is a tradeoff between increasing biofuel tolerance and achieving high biofuel yield (sustained microbial growth). Most importantly, the reaction rates in the system are uncertain and it is difficult to fine tune the system.

The ODE model of this example is reported in SI Section 4.3 (nominal parameters are in Table S4). Fig. 5E, right, shows a schematic of the microbial biofuel production process: the biofuel production pathway is represented in green, and the efflux pump expression pathway is represented in purple. We model a closed-loop system in which the Toehold Switch Brink Controller is used to force the biofuel concentration *Y* to track a given reference concentration, and we compare it to an open-loop system in which the production of *Y* is simply tuned with an inducer. In SI Section 4.3 we provide a thorough analysis of the equilibrium maps of the process, and we show that the closed-loop system presents a single equilibrium.

As done in the earlier examples, we varied the nominal parameters of the process by randomly perturbing them in the interval *±* 20% their nominal value: Fig. 5E1 shows that the closed loop system tracks linearly the reference at steady-state. Poor performance occurs when the equilibrium falls in the saturated region of the controller (inset plot), but the controller handles well the majority of random parameter combinations (Fig. 5E2). Finally, the example simulation in Fig. 5E3 shows that the controller performance can be compromised in the presence of production of *U* ^*\**^, with rate constant *ρ*_*c*_, and degradation of both *U \** and *U*, with rate constant *δ*_*c*_. We show that this issue is mitigated if *δ*_*c*_ is substantially slower than the controller degradation rate parameter *ϕ*_*c*_ (degradation of *Y*_*R*_ and *R*_*R*_); *ρ*_*c*_ = 0.5*δ*_*c*_ to guarantee an equilibrium value *u*^*tot*^ *≈* 500 *nM*. Further, consistently with expression (14), it is helpful if production rate constants *κ*_*c*_ and *θ*_*c*_ are larger than the nominal value (Table S4). Additional simulations showing the sensitivity of the closed loop system to variations of individual process parameters are in SI Section 4.3.

## 4 Discussion and conclusions

Feedback controllers in synthetic biology have the potential to mitigate many challenges that limit the performance and scalability of nonlinear molecular circuits, such as uncertainty, sensitivity to disturbances, and lack of modularity [77]. We have described general design principles to build molecular controllers for closed-loop reference tracking, disturbance rejection, and robust operation despite uncertainty in the process parameters. These principles concern the stationary input-output map of the controller, which should be ultrasensitive and have a tunable threshold that can be set by an external reference; these features naturally force the unique closed-loop equilibrium of the system to operate near the reference in a robust manner. Stability of this general architecture for closed-loop control can be assessed with linearization-based methods.

We can draw a parallel between ultrasensitive modules in a biological feedback loop and high-gain feedback controllers in engineering. High-gain negative feedback is known to improve the reliability of systems that are uncertain: operational amplifiers are a classical example that illustrate how the output of a high-gain device can precisely track the input, despite uncertainty in the gain, as long as negative feedback is present. Potential disadvantages of high-gain controllers include instability (which in our case could introduce oscillations), and chattering, a phenomenon where the controller rapidly switches between its on and off states and often occurs in sliding mode controllers [78]. Instability can be prevented by imposing suitable limitations on the closed-loop gain; if the process gain is fixed, this gain reduction would fall on the controller and possibly reduce its ultrasensitivity. This issue points to a tradeoff between stability and robustness in this type of control architecture. The problem of chattering is not expected to occur in our setup. This is because our controller is required to exhibit ultrasensitivity at steady-state, rather than instantaneously switching between its minimum and maximum output. For this reason we do not observe chattering in our computational analysis, even though the controller converges to steady-state faster than the process in the particular examples we consider.

Many natural biological sensors and controllers rely on ultrasensitivity to operate. Allosteric binding of proteins, which is known to be ultrasensitive, can be used to generate logarithmic sensors [79]. The yeast osmoregulation system combines ultrasensitivity of the MAPK pathway with negative feedback to achieve perfect adaptation [80, 81, 78]. Adaptation has also been observed in ultrasensitive enzymatic networks examined using *in silico* evolutionary algorithms [82]. Our results confirm the importance of ultrasensitivity as we point out that it confers robustness to equilibria, which in turn can be determined by the response threshold. Although we have not investigated the properties of ultrasensitive feedback control in a stochastic setting, it has been demonstrated that a steep Hill-type controller reduces the variance of protein concentration in cellular populations [83].

Ultrasensitive controllers could be built with a variety of components (see overview in Box 1), however few approaches allow rational design of both the gain and threshold. By combining sequestration reactions (comparator) and an activation/deactivation cycle (switch) [84, 33] we were able to identify an ultrasensitive network, the Brink motif, whose gain and threshold can be tuned individually: the gain is determined by specific reaction rates, and the threshold is set by the concentration of one of its inputs. According to our computational analysis, ultrasensitivity is achieved in a range of biologically plausible parameters (Fig. 3 and Table 1). As several other reaction networks for closed-loop control [15, 16, 14, 18], the Brink motif relies on sequestration, which is expected to yield a high input-output gain. There is evidence that this mechanism can also produce integral action [11, 12], however only in the ideal condition of fast sequestration and negligible degradation/dilution rates [19]; the latter requirement may be difficult to achieve in living cells. Ultrasensitivity of the Brink controller guarantees a small steady-state error which becomes zero in the limit of fast sequestration and switching rates, so the motif can operate as an integral controller. The motif can be adapted to have a monotonically increasing or decreasing ultrasensitive response, by fixing the concentration of one of its inputs. The input-output behavior of the controller should “counteract” the input-output behavior of the process, so their steady-state maps intersect in a single point and their interconnection creates a negative feedback loop. This motif could be adapted to build feedback dynamic circuits such as oscillators and bistable networks [85, 32].

The Brink controller could be experimentally realized with a variety of mechanisms. We suggest two implementation routes that rely on RNA molecules to perform mutual sequestration and regulation of the output *U*. The kinetics of RNA production and degradation, and the achievable sequestration rates make it possible to naturally build the comparator stage of the motif. According to recent *in vitro* experiments, the speed of aptamers [52] regulators for viral RNA polymerases are sufficient to operate as switches and generate ultrasensitivity; while the kinetics of toehold switches [74] have not yet been characterized *in vivo*, rates of strand displacement on similar systems *in vitro* approach those required for good performance of the controller [76]. The rapidly expanding toolkit of regulatory components based on CRISPR/Cas system offers other avenues to build the switch portion of the Brink motif [85]. As RNA-based synthetic biology is rapidly expanding [60], this motif has the potential to be a versatile and tunable element to build controllers and other dynamic devices.

## Acknowledgement

We thank Yili Qian, Harrison Steel, Franco Blanchini, and Mustafa Khammash for useful comments and discussions. This work has been supported by the National Science Foundation through grant CMMI-1266402; by the Defense Advanced Research Projects Agency through contract HR0011-16-C-01-34; and by the U.S. Department of Energy, Office of Science, Office of Basic Energy Sciences, under Award Number DESC0010595, which partially supported salary to EF and to CC.

